# From ancient fears to airborne threats: fMRI insights into neural fear responses

**DOI:** 10.1101/2025.02.10.636842

**Authors:** Silvie Rádlová, Anna Pidnebesna, Aleksandra Chomik, David Tomeček, Jaroslav Hlinka, Daniel Frynta, Eva Landová

## Abstract

Threat perception is a fundamental aspect of human cognition, shaped by evolutionary pressures and modern environmental demands. While ancestral threats (e.g., snakes) have been shown to elicit stronger neural responses than modern threats (e.g., guns), less is known about how the brain processes airborne threats, such as depictions of individuals wearing face masks. This fMRI study investigates neural responses to ancestral, modern, and airborne threats to identify shared and distinct activation patterns.

Sixty participants viewed visual stimuli from the three categories while undergoing fMRI scanning. Results showed heightened activation in the fear-processing network for all affective stimuli. In addition, activation of the ventral attention network was found for the ancestral threats. Modern threats elicited less intense responses, primarily engaging cortical regions associated with context-specific analysis. Notably, airborne threats elicited neural responses of similar intensity to ancestral threats but activated cortical regions overlapping with those for modern threats. This dual pattern highlights the brain’s capacity to integrate evolutionary biases with socially constructed threat awareness. These findings underscore the importance of recognizing airborne threats as a unique category of threat processing, with implications for public health and mental well-being.

## Introduction

The study of fear has long captivated scientists, driven by the understanding that fear is a fundamental emotion critical for survival. Fear responses can be traced back to ancient evolutionary pressures, with distinctions made between ancestral (phylogenetic) threats, which our ancestors faced, and modern (ontogenetic) threats, which have emerged relatively recently (Bennett, 2019). The aim of this study is to investigate the differences between ancestral and modern threats, including the recent threat of airborne diseases, in relation to the emotion of fear and the neural substrates involved in processing this emotion.

### Fear module and biological preparedness theory

Two pivotal theories that have shaped this area of study are the biological preparedness theory proposed by Seligman (1971) and the fear module theory developed by Öhman & Mineka (2001). Seligman’s theory of biological preparedness emphasizes the evolutionary origins of phobias and suggests that our fear responses are not random but rather shaped by our evolutionary history, where organisms that quickly learned to fear and avoid certain dangers were more likely to survive and reproduce. In theory, fears of stimuli such as spiders, snakes, or angry faces, are thought to be easier to acquire and more resistant to extinction compared to fears of stimuli that were less likely to pose immediate threats to survival (reviewed in McNally, 2016, but see also Åhs, Rosén, Kastrati, Fredrikson, Agren & Lundström, 2018 and Del Giudice, 2021 for critical evidence). Generally, it was supposed that these stimuli, referred to as phylogenetic threats, were inherently more salient and evoked stronger and more persistent fear responses compared to neutral stimuli and even to modern, ontogenetic threats like guns and electrical outlets.

Building on the concept of biological preparedness, Öhman & Mineka (2001) introduced the fear module theory, which posits the existence of a specialized neural circuitry in the brain dedicated to rapid detection and response to fear-relevant stimuli. This fear module is believed to be automatic, unconscious, and specifically tuned to ancestral threats. The theory proposes that this module operates independently of higher cognitive processes, allowing for swift and efficient responses to dangers that were common in the environment of evolutionary adaptedness (Öhman & Mineka, 2001). This neural mechanism is thought to prioritize survival by enabling quick reactions to life-threatening situations. The current neural and behavioral model of fear (commonly referred to as the survival circuit) conceptualizes fear as the organism’s complex response to life-threatening and other situations. This model encompasses both automatic fear and defensive reactions, as well as learned fear and its cognitive regulation, including the prediction of danger and the formulation of planned responses (Mobbs, Headley, Ding & Dayan, 2020). Within this framework, the immediacy of the threat and the stage of the organism’s response play significant roles. Nonetheless, the fear module proposed by Öhman & Mineka (2001) offers a valuable framework for understanding non-associative, evolutionarily rooted fears, such as the fear of snakes (Kawai & Kawai, 2019).

### Phylogenetic vs ontogenetic threats

Numerous studies have relied on these theoretical frameworks to compare responses to phylogenetic and ontogenetic threats. These studies typically examine the resistance to extinction and physiological responses to different types of threats (McNally, 1987; see Del Giudice 2021 for a metanalysis). Research has shown that in some cases, fear responses to phylogenetic threats are more robust and persistent compared to those elicited by ontogenetic threats. For example, participants exhibit greater resistance to the extinction of conditioned fear responses to snakes and spiders compared to modern threats like guns and syringes (Hugdahl & Kärker, 1981; Cook, Hodes, & Lang, 1986; Öhman & Mineka, 2001).

In a study by Tomarken, Sutton & Mineka (1995), only phylogenetic fear-relevant stimuli, as opposed to ontogenetic ones (e.g., slides depicting damaged and exposed electric outlets), were found to be associated with covariation biases for shock. In a task designed to assess covariation or illusory bias, participants were shown slides of stimuli categorized as either fear-relevant (e.g., snakes) or fear-irrelevant (e.g., flowers). After each slide presentation, participants experienced one of three outcomes: a shock, a tone, or no further stimulus. They were instructed to estimate the correlation between each slide type and its respective outcome. The findings revealed that participants tended to overestimate the magnitude of the association between fear-relevant stimuli and shock when the stimuli were snakes.

Similarly, the influence of angry faces (a widely studied phylogenetic threat) on attention has been demonstrated in a seminal study (Öhman, Lundqvist & Esteves, 2001). Using a visual search task that measured the speed of detecting a target stimulus within a distractor matrix, the researchers showed that threatening faces were identified more quickly than happy or sad faces. Additionally, snakes, identified as a potential phylogenetic threat (Isbell 2006, 2009), were found to be detected faster than other animals in visual search tasks employing eye-tracking technology (Shibasaki & Kawai, 2011; Soares, Lindström, Esteves & Öhman, 2014). Recent evidence, nonetheless, indicates that the superiority effect of threatening stimuli on attention is not limited to ontogenetic or phylogenetic threats but may represent a more generalized phenomenon (reviewed in Zsido, Deak & Bernath, 2019).

However, the empirical support for these theories is not without challenges. The behavioural outcome of many studies, typically focusing on fear conditioning, illusory correlation, or reaction time, found limited evidence supporting the notion that ancestral fears are universally superior to the modern ones (see review by Shapouri & Martin, 2022, and also Abado, Aue & Okon-Singer, 2023, Zsido, Hout, Hrrnandez, White, Polák, Kiss & Godwin 2024). This discrepancy may be attributed to various methodological issues (e.g., usage of ambiguous stimuli or varying context, Cao et al., 2014, see discussion), but it can also reflect different effects of ancestral stimuli to human behavior than what the researches have presumed. What if the resistance to extinction or reaction time are not the factors defining the distinction between the ancestral and modern threats? The observed discrepancy may be attributable to differences in neural activation patterns. Visual stimuli showing ancestrally relevant threats may activate quick, automatic, subcortical, unconscious pathways of fear processing (as demonstrated in studies using event-related potentials (ERPs) by Zhang and Guo, 2020, and further reviewed in the context of fast and unconscious fear processing of snakes by Wang, Cong & Hu, 2023; Biondi, Gomes, Maior & Soares, 2024; and Setogawa, Matsumoto, Nishijo & Nishimaru, 2024; or may trigger more intense activation of the fear network, see below). While the first mentioned case cannot be easily studied using functional magnetic resonance because its response is slow, the second case was successfully shown by, i.e., Dhum, Herwig, Opialla, Siegrist & Brühl, 2017). These authors have demonstrated that evolutionarily threatening pictures evoked significantly stronger activations than modern-threatening pictures in most regions of the network for processing threatening stimuli (the left inferior frontal gyrus, right middle frontal gyrus, right parietal lobe, right precuneus, left thalamus, bilateral fusiform gyrus, bilateral superior parietal lobule, and bilateral amygdala), although the respondents consciously ranked the modern stimuli as more threatening. Similarly, Fang, Li, Chen & Yang, (2016) found stronger activation of the amygdala and subcortical regions in response to evolutionary threats (snakes), especially in a human context.

### The Fear Network in the Human Brain

Recent fMRI studies suggest that emotional categories, including fear, are uniquely represented within brain activity. However, these systems are highly distributed, involving contributions from both subcortical regions (e.g., thalamus, periaqueductal gray, basal forebrain, and amygdala) and cortical regions (e.g., prefrontal cortex, anterior and midcingulate cortex, and insular cortex; Zhou et al., 2021; Kragel & LaBar, 2016). Additionally, the cerebellum has been implicated in these processes (Tao, He, Lin, Liu & Tao, 2021). Despite this widespread distribution, there is compelling evidence highlighting the critical roles of specific regions in neural threat processing.

The traditionally recognized central structure of the fear network is the amygdala (Phelps & LeDoux, 2005; Vuilleumier, 2005; Pourtois, Schettino & Vuilleumier, 2013). It is crucial for detecting and responding to threat-related stimuli. It processes fear rapidly, even outside of conscious awareness, activating in response to fear-relevant images and sounds, as seen in both animals and humans (Sander, Grafman & Zalla, 2003; Méndez-Bértolo et al., 2016; Silva, Gross & Gräff, 2016). This rapid response mechanism is thought to arise from a specialized subcortical pathway that bypasses the cortex, allowing the amygdala to quickly assess potential dangers and initiate defensive behaviors (Tamietto & De Gelder, 2010; Tovote, Fadok & Lüthi, 2015; Morris, Öhman & Dolan, 1999; Wang, Luo, Chen, Luan, Wang, Wang & Fang, 2023); This subcortical pathway connects visual inputs to the amygdala through the superior colliculus and pulvinar (McFadyen, Mattingley & Garrido, 2019), allowing rapid emotional processing of visual threat cues like fearful faces, even before full conscious awareness. This pathway supports quick, automatic responses to potential threats without cortical involvement, essential for survival in rapidly changing environments (Bertini & Làdavas, 2021).

An additional subcortical brain area directly connected to the amygdala is the nucleus accumbens (Tamietto & De Gelder, 2010), which plays a role in integrating affective and motivational signals from both the amygdala and prefrontal cortex. Its dopaminergic circuits are engaged in response to rewarding or threat-related cues, which can promote approach or avoidance behaviors as needed (Floresco, 2015; Salamone, 1994). These interconnected areas form a highly efficient system for detecting, processing, and responding to fear, supporting the theory of a specialized neural circuitry for handling fear-relevant stimuli.

Major cortical areas involved in the processing of fear are the prefrontal cortex (PFC) and anterior cingulate cortex (ACC). The PFC plays a regulatory role in processing and modulating the amygdala’s activity, allowing for cognitive control over fear responses (Das et al., 2005). Specifically, the ventromedial PFC (vmPFC) aids in fear extinction by inhibiting amygdala responses, thus helping to moderate reactions to perceived threats (Sotres-Bayon & Quirk, 2010; Lesting, Narayanan, Kluge, Sangha, Seidenbecher & Pape, 2011). The ACC monitors emotional conflicts and integrates sensory and emotional information, modulating the intensity of fear responses. It works in coordination with the amygdala and PFC, particularly during high-stakes decisions or emotionally challenging tasks (Das et al., 2005; Hofmann, Ellard & Siegle, 2012; Lindner, Neubert, Pfannmöller, Lotze, Hamm & Wendt, 2015).

The brain also plays a crucial role in the cognitive assessment of imminent danger. In experiments simulating immediate threats, activation was observed in the midbrain periaqueductal gray (PAG) and the medial cingulate cortex (MCC). Conversely, when a threat is perceived as distant or when the situation is ambiguous, allowing time for decision-making, circuits associated with cognitive evaluation become more active. These include the posterior cingulate cortex (PCC), the hippocampus (HIPP), and the ventromedial prefrontal cortex (vmPFC). These regions support complex information processing, the development of cognitive strategies for threat avoidance, and overall behavioral flexibility (Qi, Hassabis, Sun, Guo, Daw & Mobbs, 2018).

The amygdala and the orbitofrontal cortex also help modulate sensory processing by enhancing the responsiveness of visual areas (Tabbert, Stark, Kirsch & Vaitl, 2005), thereby prioritizing emotionally relevant stimuli over neutral ones (Junghöfer, Bradley, Elbert & Lang, 2001; Bradley, Sabatinelli, Lang, Fitzsimmons, King & Desai, 2003). Studies utilizing functional magnetic resonance imaging (fMRI) and positron emission tomography (PET) have shown that exposure to emotionally salient visual stimuli leads to increased neural activity in areas like the fusiform gyrus, occipital cortex, and inferotemporal regions, which are responsible for processing complex visual information (Lang, Bradley, Fitzsimmons, Cuthbert, Scott, Moulder & Nangia, 1998). In our previous study of fear of spiders on arachnophobic respondents, we confirmed these results (Landová et al, 2023).

The insula, particularly its anterior regions, is involved in representing interoceptive states and conscious emotional awareness, serving as a hub for integrating bodily states with emotional experiences. This makes the insula crucial for processing a wide range of emotions (including fear), as it enables individuals to feel and interpret bodily sensations associated with different emotions (Herbert, Herbert & Pauli, 2011; Suzuki, 2012). However, the insula’s response to emotional stimuli seems to unfold in stages. Initial activation in response to emotionally salient stimuli occurs around 200 ms in the right insula, which is non-specific to emotion type, followed by a more differentiated response for disgust, around 350 ms. (Chen, Dammers, Boers, Leiberg, Edgar, Roberts & Mathiak, 2009; Krolak-Salmon et al., 2003).

### Airborne diseases and the COVID-19 pandemic: modern threats

Recently, we faced a threat that is inherently invisible—the COVID-19 pandemic. Airborne transmission via respiratory droplets and aerosols (Wang, Prather, Sznitman, Jimenez, Lakdawala, Tufekci & Marr, 2021) is heavily influenced by human mobility, frequency of contact, and population density (Kucharski et al., 2020). For most of human evolutionary history, small, isolated groups limited inter-group interactions, resulting in localized epidemics (Weisdorf, 2005). This dynamic shifted approximately 10,000 years ago with the emergence of cities and extensive trade networks (Cockburn, 1971). The subsequent increase in population density enabled the rapid spread of airborne pathogens (Kakehashi & Yoshinaga, 1992). From an evolutionary standpoint, the threat of airborne diseases that can escalate to pandemic levels is predominantly a modern phenomenon (Moreno & Gibbons, 2022; Peléšková, Polák, Janovcová, Chomik, Sedláčková, Frynta & Landová, 2024; but see Silva, Moura, Ferreira Júnior, Nascimento & Albuquerque, 2022).

During the COVID-19 pandemic, individuals often remained asymptomatic in the early stages of infection (Furukawa, Brooks & Sobel, 2020). However, visible symptoms such as coughing, fever, and fatigue—associated with the later stages of viral infection—align with cues that the behavioral immune system (BIS) has evolved to detect since ancient times (reviewed in Thiebaut, Méot, Witt, Prokop & Bonin, 2021). During the pandemic, however, we frequently encountered symbolic, indirect cues of viral presence: individuals wearing masks, hand-sanitizer stations in public spaces, and images of hospitals on television. These indirect cues can thus be considered “modern threats,” as masks and hospitals were absent in early human history when the BIS (Curtis, Aunger & Rabie, 2004; Schaller and Park, 2011) developed (Seitz et al., 2020). Individual differences in BIS reactivity, including germ aversion and pathogen-related disgust sensitivity, were shown to predict concerns about COVID-19 and adherence to preventive health behaviors (Shook, Sevi, Lee, Oosterhoff & Fitzgerald, 2020). These findings highlight how an evolutionarily ancient system, originally adapted to avoid contamination, can respond to modern airborne pandemics (such as the novel coronavirus Covid-19).

The current study seeks to examine how these relationships manifest in neural fear processing of threats posed by airborne pandemics. Specifically, we investigate whether stimuli representing ancestral threats through modern cues activate the BIS and elicit neural activity patterns similar to those triggered by traditional ancestral fears. This is one of the core questions our research aims to address.

### General aims

Our study is divided into two parts. The first part examines neural-level differences in fear responses to ancestral versus modern threats. The second part compares ancestral and modern stimuli with those depicting modern airborne disease cues, aiming to determine whether the response aligns more closely with ancestral or modern threats. More specifically, we formed the following research questions and hypotheses: (1) the “pure emotions” evoked by photographs of threatening situations according to the tested conditions (ancient fear, modern fear, and airborne diseases) would result in significant activation of the fear network. To investigate this, we examined their contrasts with scrambled control images. We also anticipated that (2) the neural activation within the fear network would be more pronounced during the presentation of stimuli depicting ancestral threats compared to modern threats. Finally, (3) considering the ancient origins of the behavioral immune system involved in the perception of airborne disease threats, we hypothesized that neural responses to such threats would parallel those elicited by ancestral fears. To test this, we directly compared neural activation patterns associated with airborne disease threats to those evoked by both ancestral and modern fear stimuli. Additionally, we employed complementary methods to analyze the intensity and distribution of neural responses relative to these categories of threats.

## Methods

### Participants

We recruited a total of 60 healthy women aged between 19-47: 47 were of the 19-35 category, 17 were older (mean = 29.57; median = 27). Fifty of them were of Czech nationality, 10 of other nationalities (7 different nationalities). Twenty respondents had biological education, 25 social education, and nine had other types of education.

The participants completed a series of questionnaires (see below) during the recruitment phase and were subsequently invited to participate in the fMRI experiment. They were thoroughly briefed on the experimental procedures and provided written informed consent. Participants were instructed that they would be shown photographs depicting various types of threats and were asked to imagine these threats as personally relevant. Following this preparation, they underwent the fMRI session, during which their neural responses to the stimuli were recorded. To ensure the stimuli used in the study evoke the predicted emotion (fear), the respondents also evaluated all stimuli included in the experiment on a 7-point scale according to their perceived fear and disgust (1 corresponded to low values, 7 to high)

At the conclusion of the session, participants took part in a debriefing with the experimenter to ensure their well-being and to address any concerns. They were also offered additional information about the experiment and given the opportunity to ask questions for clarification.

All of the respondents filled in SNAQ-12 (only three respondents, i.e., 5% of the respondents[KV1], scored 8 or higher; the minimum score was 0, the maximum was 11, the mean of the scores was 3.2, the median was 3) and FCV19-S (13 respondents scored 16 or higher, i.e., 21.7% respondents were likely to experience high fear of the COVID disease; the minimum score was 7, maximum was 28, mean of the scores was 11.6, median was 9.5). These values indicate that the participants in the study corresponded to the general population and did not exhibit significant hypersensitivity either to the fear of snakes or to viral diseases such as COVID-19.

### Ethics statement

All procedures performed in this study were conducted in accordance with the local legislation and institutional requirements. The studies involving humans were approved by the Ethics Committee of National Institute of Mental Health National Institute of Mental Health (no. 91/21, granted 31 March 2021). All the experiments were carried out following the 1964 Helsinki declaration and its later amendments or comparable ethical standards. Recruitment of the respondents for this study has started on 9^th^ August, 2023 and lasted till 21^st^ February, 2024. Written informed consent was obtained from all participants included in the study.

### Experimental design

#### Stimuli preparation

We prepared 60 visual stimuli (30 unique photos and 30 horizontally transferred copies) depicting ancestral fear (condition: AF60), 60 depicting modern fear (condition: MF60), and 60 depicting airborne diseases (condition: DIS60; see Figure 1). The pictures were selected either from existing emotional picture databases (IAPS, Lang, Bradley & Cuthbert, 2005; SMID, Crone, Bode, Murawski & Laham, 2018; EmoMadrid, Carretié, Tapia, López-Martín, Albert, 2019), Internet resources such as Pixabay (www.pixabay.com), or photographed by us. The snake pictures were used from a previous fMRI study (Landová et al. 2023). Rarely, we used photos from Czech Internet newsletter magazines (Aktuálně.cz, Hospodářské noviny and Lidovky; see S1 Supplement). We selected only pictures that were of a high quality and emotional valence (according to the databases or our previous experiments). Each picture was re-scaled to 1024 x 768 pixels for optimal fMRI presentation. Also, because by character, these stimuli are visually very different and incomparable, each condition has had its own control that consisted of the same 60 images, but scrambled: MF60sc, AF60sc, DIS60sc; These pictures were generated using a special software we developed that allowed for mass-conversion of pictures into their scrambled versions. Each picture was sliced into a specific number of pieces (in this case, 31 x 31 pieces, i.e., 961 pieces total), which were then assembled together in a random order to create a new, scrambled stimulus. Scrambled images retained the same visual properties (lightness, contrast, hue) as the original images, but lacked the emotional charge of each image because the objects became unidentifiable (see Figure 2). In total, we had six conditions (AF, MF, DIS, and their scrambled copies) consisting of 360 images (60 per condition).

**Figure 1.**
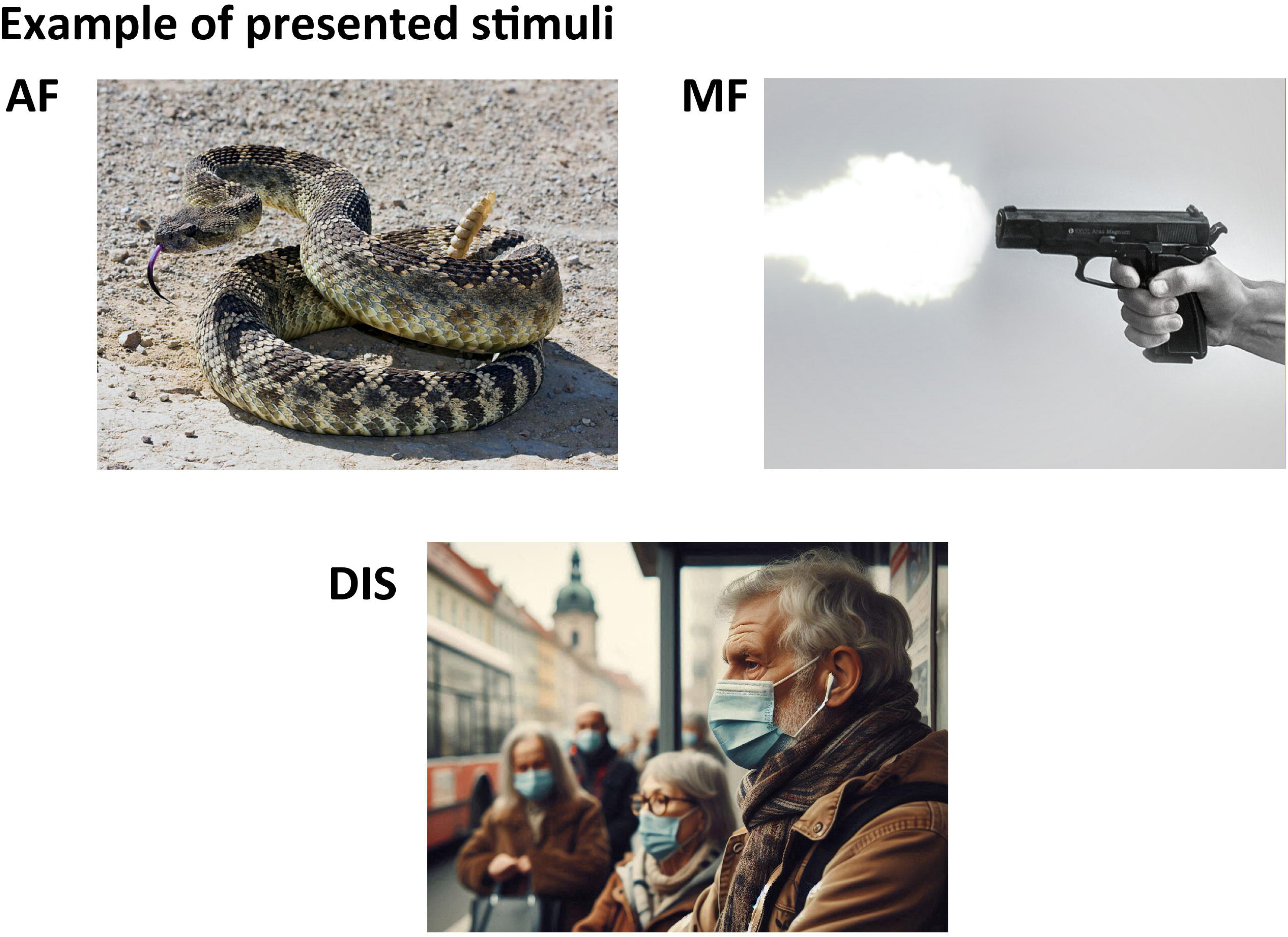
Examples of stimuli used in the study. The study included three categories of stimuli, each containing 60 images: ancestral fear (AF), modern fear (MF), and airborne diseases (DIS). The pictured AF and MF stimuli were obtained from Pixabay, shared under the “Content License”, which allows free use, and are the exact files that were used in the study; all of the **DIS** images included people in the photos; thus, to comply with BioRxiv’s policy to avoid the inclusion of photographs and any other identifying information of people, this example picture was AI-generated. It is just an example of how the stimuli looked like and was not used in the study.

**Figure 2.**
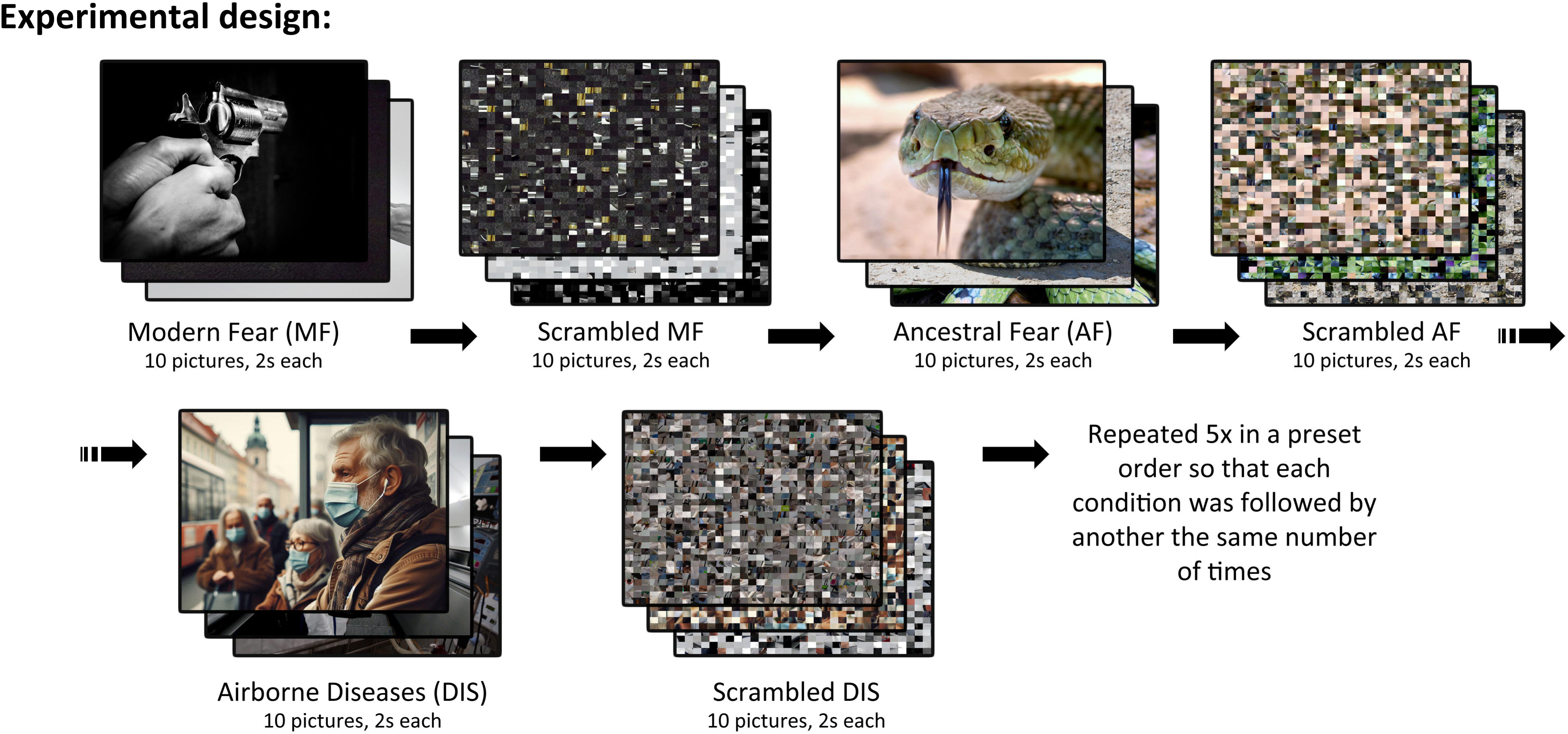
Schema of the Experimental Design. Modern and ancestral fear-relevant stimuli were presented alongside images depicting airborne disease-related scenes. Each experimental condition was followed by a block of scrambled control images. Stimuli were presented in blocks lasting 2 seconds each to a total of 60 respondents. The example photos of the soldier and the snake are from Pixabay, shared under the “Content License”, allowing free use. All stimuli depicting airborne diseases in this study featured individuals. The image shown here is an example of the type of stimuli used, but this specific photo was not part of the study. In accordance with BioRxiv’s policy to avoid including photographs or any identifying information of individuals, we replaced the original image with an AI-generated example.

All pictures were scored by the respondents on a 7-point Likert scale according to perceived fear and disgust, and Tukey test confirmed significant differences between the scoring of fear and disgust of each condition. See Table 1 for a summary of means and medians of the scores for each condition.

**Table 1.** Mean and median scores of the emotional stimuli presented in Experiment 2 as scored by the participating respondents. All pictures were scored on a 7-point Likert scale based on self-reported fear and disgust. Tukey test confirmed a significant difference in scoring of fear and disgust in each condition, and each condition elicited fear significantly more than disgust.

|  | Fear |  | Disgust |  | p |
| --- | --- | --- | --- | --- | --- |
|  | Mean | Median | Mean | Median |  |
| Modern Fear (guns) | 4.32 | 5 | 2.68 | 2 | <0.001 |
| Ancestral Fear (snakes) | 4.20 | 5 | 2.61 | 2 | <0.001 |
| Airborne Diseases | 2.99 | 2 | 2.29 | 1 | <0.001 |

#### Stimuli presentation

During the fMRI experiment, we presented 10 images from each condition in a block lasting 20 seconds (i.e., one stimulus was presented for 2 s). After each block of emotional condition, its specific control followed (for example, after MF60, MF60sc always followed, including scrambled versions of the same set of pictures n a randomized order; i.e., if the first block contained pictures 01, 50, 23, 13, 59, 07, 29, 28, 35, and 18, the following control showed the same scrambled pictures but in a different order, such as 29sc, 35sc, 07sc, 01sc, 28sc, 59sc, 23sc, 18sc, 13sc, and 50sc). The order of presentation of the emotional condition blocks was preset in a pseudo-random order so that each block was followed by another the same number of times (MF60, AF60, DIS60, AF60, DIS60, MF60, DIS60, AF60, MF60, AF60, MF60, DIS60, MF60, DIS60, AF60, DIS60, MF60, AF60). The whole fMRI measurement lasted nine minutes.

#### Measurements

The MRI data acquisition was performed on a 3 Tesla Siemens Prisma scanner equipped with a standard 64-channel. Structural 3-dimensional (3D) images were obtained for anatomical reference using the T1-weighted (T1w) magnetization-prepared rapid gradient echo (MPRAGE) sequence with the following parameters: repetition time (TR) of 2100ms, echo time (TE) 2.3ms, flip angle 8°, voxel size of 0.9×0.9×0.9mm3, field of view (FOV) 288mm×288mm, matrix size 320×320, 224 sagittal slices.

All functional images were obtained using the T2*-weighted (T2*w) gradient echo-planar imaging (GR-EPI) sequence sensitive to the blood oxygenation level-dependent (BOLD) signal with the following parameters: repetition time (TR) of 1000ms, echo time (TE) 30ms, flip angle 52°, voxel size of 3×3×3mm3, field of view (FOV) 222mm×222mm, matrix size 74×74, each volume with 68 axial slices (multiband sequence with factor 4). In total, 720 volumes were acquired during the measurement.

During the preprocessing, the structural and functional images were converted from DICOM to NIFTI format using the dcm2niix tool (Li, Morgan, Ashburner, Smith & Rorden, 2016). The functional images were then corrected for intensity variations caused by the bias field. In the next step, we used a default preprocessing pipeline for volume-based analyses in the CONN toolbox (Whitfield-Gabrieli & Nieto-Castanon, 2012), which performs the following steps with the functional data: functional realignment and unwarp; slice-timing correction; outlier identification; direct segmentation and normalisation to the MNI (The Montreal Neurological Institute) space; and finally, spatial smoothing of the functional data with 8mm FWHM (full width at half maximum) kernel. The spatially smoothed data were then used for the following statistical analysis. As the final step in data preparation, we conducted quality control, which resulted in the exclusion of 6 out of 60 subjects due to excessive head motion.

## Statistical Analysis

As was mentioned before, in the course of our analysis, the following hypotheses were evaluated:

1. **“Pure” emotional reaction:** We hypothesised that exposure to fear-evoking stimuli and stimuli showing airborne diseases, compared to their scrambled controls (i.e., “pure” emotion condition), would result in significant activation of the fear network.
2. **Ancestral vs. Modern Stimuli:** We anticipated that neural activation of the fear network would be stronger during the presentation of ancestral stimuli compared to modern stimuli.
3. **Airborne Disease vs. Ancestral vs. Modern Fear Stimuli:** We predicted that the average neural response to stimuli depicting airborne diseases would be of similar intensity as the response elicited by ancestral stimuli. Furthermore, we also aimed to investigate the activation profiles associated with disease-related stimuli, as well as ancestral and modern stimuli, to determine whether the neural response to airborne diseases more closely resembles the response to modern or ancestral stimuli.

Hypothesis testing was performed using several tools. Statistical Parametric Mapping (SPM12; Wellcome Trust Centre for Neuroimaging, London, United Kingdom) and our own pipelines were used for the analysis.

In the first step, the first-level statistic was performed in SPM12. For that, we created several variables indicating stimuli of several types: ancestral fear (AF), modern fear (MF), airborne disease (DIS), and their scrambled versions (AFsc, MFsc, DISsc, respectively). Then, the general linear model (GLM) was built, and three contrasts, AF>AFsc, MF>MFsc, and DIS>DISsc, were computed per subject using a one-sample t-test.

We conducted a second-level group analysis to test **the first hypothesis**, which examines the fear network activation during the experiment. Specifically, we performed a one-sample t-test at the group level for the contrasts AF > AFsc, MF > MFsc, and DIS > DISsc. Statistical significance was assessed using a family-wise error (FWE) correction with a threshold of p < 0.05 to control for multiple comparisons.

To test **the second hypothesis**, which explores the differences in brain activation between ancestral and modern stimuli, we conducted a second-level group analysis performing a paired t-test for the contrast AF > AFsc versus MF > MFsc. We computed the contrast in both directions, [(AF > AFsc) > (MF > MFsc)] and [(MF > MFsc) > (AF > AFsc)]. As with the first hypothesis, significance was determined using a family-wise error (FWE) correction with p < 0.05 to account for multiple comparisons.

To test **the third hypothesis**, which examines the relationship between airborne disease stimuli and ancestral and modern fear stimuli, we used the first-level t-maps of AF > AFsc, MF > MFsc, and DIS > DISsc contrasts to compute the average activation index (see the detailed description below). Similarly to the previous paper, we averaged the t-maps masked by ROI defined in Landová et al. (2023) for every subject and every mentioned contrast. Following this, we compared the resulting distributions across subjects to assess differences in activation patterns. We used a paired two-sided Wilcoxon signed rank test for the null hypothesis that the difference between matched samples comes from the distribution with zero mean. We performed the described test for pairs of activation indices for Ancestral Fear vs Modern Fear, Modern Fear vs Disease, and Ancestral Fear vs Disease.

### The average activation index

In Landová et al. (2023), we presented the average activation index applied for arachnophobia under the name Spider Fear Index. For a given first-level t-map, we masked it by the predefined ROI and then computed the average value of the masked t-map. The mask was computed in Landová et al. (2023) on a group of arachnophobic respondents watching spider and beetle pictures, using a spider>beetle contrast (group level analysis, significance level 0.05, FWE correction), and included parts of the following areas, bilaterally: precentral gyrus, inferior frontal gyrus - opercular part, triangular part, and orbital part; insula, calcarine fissure and surrounding cortex, cuneus, lingual gyrus, superior occipital gyrus, middle occipital gyrus, inferior occipital gyrus, fusiform gyrus, superior parietal gyrus, Inferior parietal (including right supramarginal gyrus and right angular gyrus), and middle and inferior temporal gyrus. Moreover, the ROI included the right middle frontal gyrus, right parahippocampal gyrus, and right precuneus.

Further, in **the exploration phase**, we compared the activation maps obtained during the Disiase blocks with those obtained for Ancestral and Modern Fear blocks. For the qualitative analysis, we performed the second-level (group) analysis by computing the paired t-test for the contrasts [(AF > AFsc) > (DIS > DISsc)], [(MF > MFsc) > (DIS > DISsc)], [(DIS > DISsc) > (AF > AFsc)], and [(DIS > DISsc) > (MF > MFsc)] (FWE-corrected, p < 0.05).

For the quantitative analysis, the Partial Least Squares (PLS) classification was used in the following way. In this analysis stage, we worked with the first-level t-maps for the contrasts AF > AFsc, MF > MFsc, and DIS > DISsc. First, we masked all the activation maps with the average grey matter mask obtained during the data preprocessing. Then, we fitted the PLS model on the Ancestral and Modern Fear maps as regressors and the type of map (Ancestral or Modern) as a response variable. Then, the obtained model was used to predict the type of activation maps from the Disease block. In this way, we define if the neural activation during the Disease stimuli is closer to Modern or Ancestral Fear blocks. On the technical side, the number of PLS components was defined such that the explained variance for both regressors and response reached at least 90%, which resulted in 29 components. The quality of the obtained PLS model was tested using the leave-one-out approach on the set of AF > AFsc and MF > MFsc maps.

## Results

### Hypothesis 1: “pure” emotional reaction

All affective stimuli (AF > AFsc, MF > MFsc, and DIS > DISsc) elicited significant activation in overlapping regions associated with the fear network and emotional processing (see Fig 3). These included frontal regions (inferior frontal gyrus - right opercular and left orbital parts, and medial superior frontal gyrus); limbic and subcortical areas (bilateral amygdala, hippocampus, parahippocampal gyrus, thalamus); occipital and parietal regions (bilateral calcarine cortex, cuneus, lingual gyrus, superior, middle, and inferior occipital gyri, fusiform gyrus, postcentral gyrus, superior and inferior parietal gyri, supramarginal gyrus, precuneus, and angular gyrus); and temporal regions (left superior, and bilateral middle and inferior temporal gyri; see S2 Supplement).

**Figure 3.**
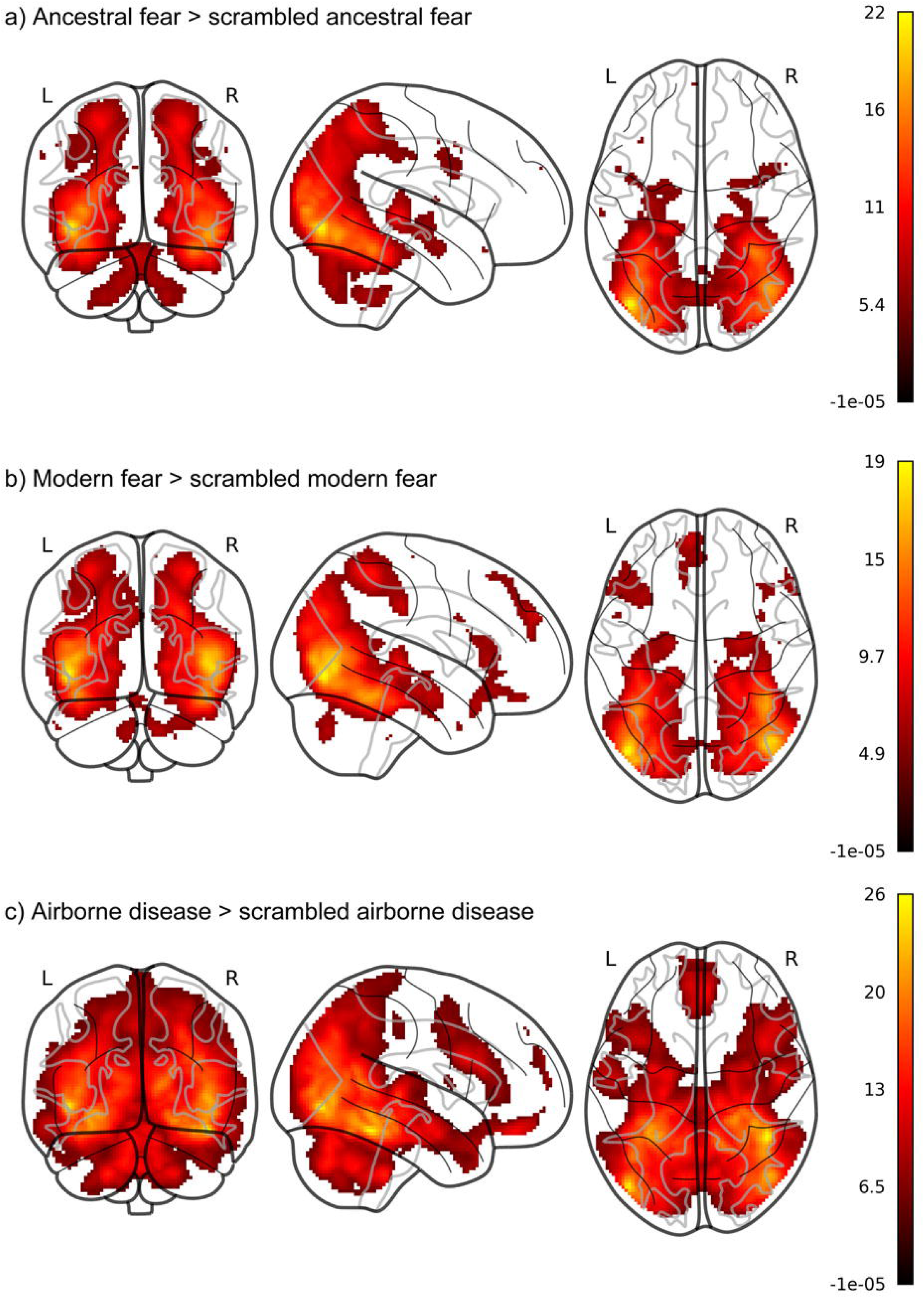
Activation map for the contrasts fear/disease > noise, thresholded at p < 0.05 (FWE corrected). The color scale represents t-statistics with low t-values shown in black and high t-values in yellow. The map highlights regions with significant neural activation differences during the **fear/disease** condition compared to **noise**. (a) Contrast ancestral fear > scrambled ancestral fear (b) Contrast modern fear > scrambled modern fear (c) Contrast airborne disease > scrambled airborne disease

The bilateral precentral gyrus, middle frontal gyrus, putamen, and pallidum were activated in both AF > AFsc and DIS > DISsc contrasts, but not the MF > MFsc. Similarly, the superior frontal gyrus, inferior frontal gyrus (triangular part), and gyrus rectus were activated in both the MF > MFsc and DIS > DISsc contrasts, but not the AF > AFsc contrast.

Moreover, there were unique activations in some of the contrasts. In the contrasts AF > AFsc, activation was observed in the left insula. Unique activations in the DIS > DISsc contrast included the paracentral lobule (bilateral), median cingulate and paracingulate gyrus (bilateral), posterior cingulate gyrus (bilateral), and middle frontal gyrus (orbital part: bilateral).

These results suggest that while all fear-related stimuli engage a common network for fear and emotional salience, the type of threat modulates specific brain regions, reflecting differences in survival relevance and processing demands.

### Hypothesis 2: Ancestral vs. Modern Stimuli

To test the hypothesis that ancestral stimuli evoke stronger activation in the fear-processing network compared to modern stimuli, we compared the activation maps for ancestral and modern fear (see Figure 4a). The contrasts revealed distinct patterns of activation with ancestral stimuli [(AF > AFsc) > (MF > MFsc)], showing heightened activation in occipital and parietal regions, consistent with enhanced visual and spatial attention to phylogenetic threats. In contrast, modern stimuli [(MF > MFsc) > (AF > AFsc)] elicited greater activation in temporal and parahippocampal areas, potentially reflecting more contextually integrated and memory-related processing of ontogenetic threats (see Table 2).

**Figure 4.**
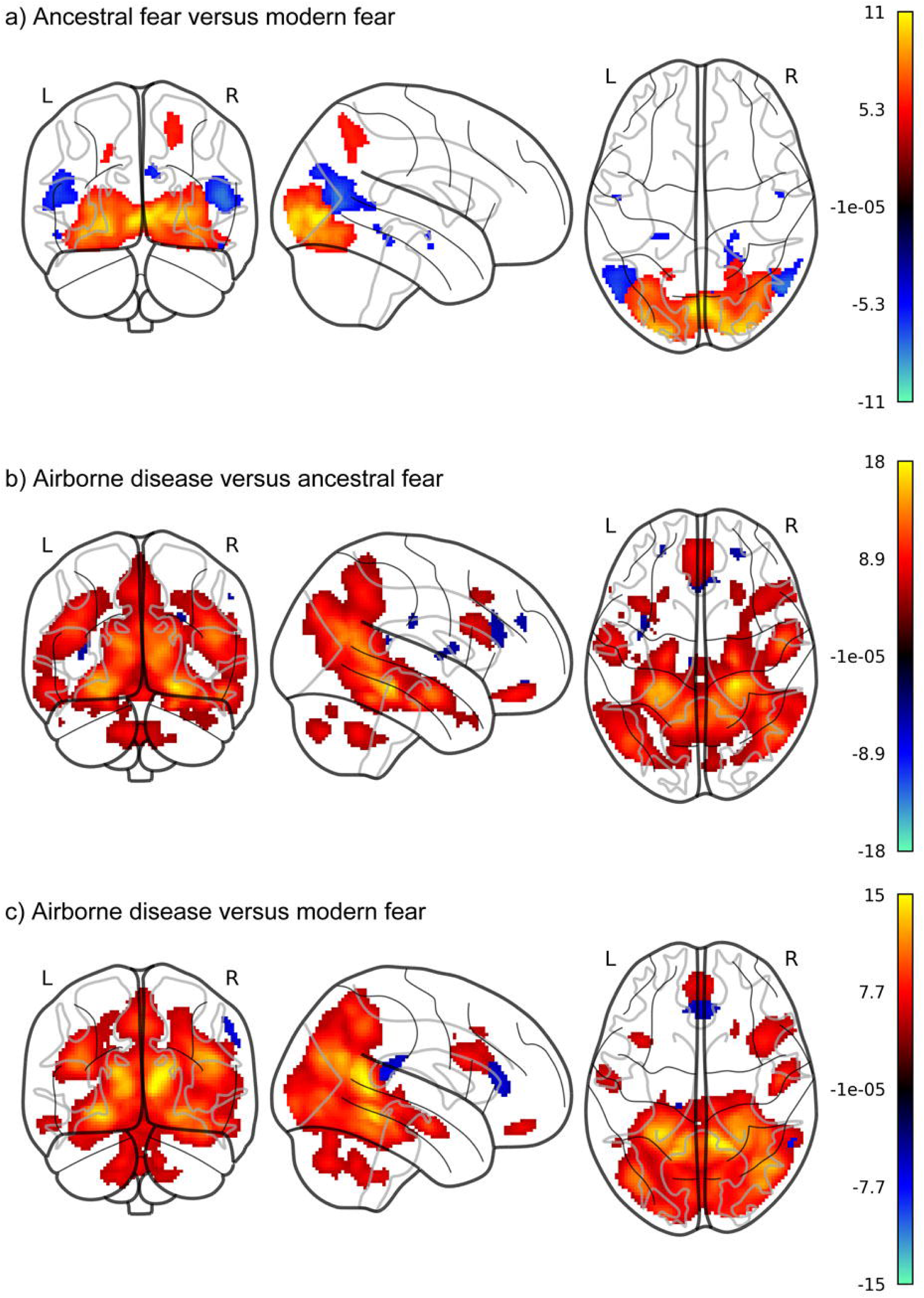
Activation map for contrasts comparing all conditions with each other. , thresholded at p < 0.05 (FWE corrected). a) The positive part of the color scale represents t-statistics for the contrast [(AF > AFsc) > (MF > MFsc)], and the negative part of the colorscale represents the contrast [(MF > MFsc) > (AF > AFsc)]. b) The positive part of the color scale represents t-statistics for the contrast [(DIS > DISsc) > (AF > AFsc)], and the negative part of the colorscale represents the contrast [(AF > AFsc) > (DIS > DISsc)]. c) The positive part of the color scale represents t-statistics for the contrast [(DIS > DISsc) > (MF > MFsc)], and the negative part of the colorscale represents the contrast [(MF > MFsc) > (DIS > DISsc)].

**Table 2.** Contrasts (AF > AFsc) > (MF > MFsc), and (MF > MFsc) > (AF > AFsc), whole-brain analysis, thresholded at cluster level p<0.05 (FWE-corrected). The presented t-scores with the corresponding MNI coordinates (x,y,z) represent the maximum values within the brain area.

**$(AF > AF_{sc}) > (MF > MF_{sc})$**
| Area |  | Extent | t-score | x | y | z |
| --- | --- | --- | --- | --- | --- | --- |
| Calcarine fissure and surrounding cortex | L | 0.188 | 9.89 | -2 | -82 | -4 |
| Calcarine fissure and surrounding cortex | R | 0.244 | 10.2 | 6 | -82 | 0 |
| Cuneus | L | 0.001 | 5.98 | -6 | -98 | 6 |
| Cuneus | R | 0.047 | 6.81 | 18 | -88 | 8 |
| Lingual gyrus | L | 0.284 | 10.3 | 2 | -80 | -2 |
| Lingual gyrus | R | 0.306 | 10.5 | 4 | -80 | -2 |
| Superior occipital gyrus | L | 0.094 | 7.12 | -10 | -94 | 4 |
| Superior occipital gyrus | R | 0.04 | 8.2 | 22 | -96 | 4 |
| Middle occipital gyrus | L | 0.173 | 9.69 | -22 | -96 | -4 |
| Middle occipital gyrus | R | 0.177 | 9.65 | 28 | -92 | 0 |
| Inferior occipital gyrus | L | 0.513 | 9.68 | -20 | -96 | -8 |
| Inferior occipital gyrus | R | 0.442 | 9.57 | 24 | -94 | -2 |
| Fusiform gyrus | L | 0.185 | 8.7 | -36 | -84 | -14 |
| Fusiform gyrus | R | 0.173 | 8.35 | 36 | -76 | -12 |
| Superior parietal gyrus | L | 0.004 | 5.34 | -18 | -60 | 42 |
| Superior parietal gyrus | R | 0.067 | 5.89 | 20 | -60 | 54 |
| Inferior parietal, excluding Supramarginal<br>gyrus and Angular gyrus | R | 0.004 | 5.14 | 26 | -52 | 50 |
| PreCuneus | R | 0.001 | 5.61 | 18 | -58 | 44 |
| Inferior temporal gyrus | R | 0.032 | 7.41 | 40 | -64 | -10 |

**(MF > MFsc) > (AF > AFsc)**
| Area |  | Extent | t-score | x | y | z |
| --- | --- | --- | --- | --- | --- | --- |
| Parahippocampal gyrus | R | 0.013 | 5.57 | 22 | -34 | -14 |
| Calcarine fissure and surrounding cortex | R | 0.015 | 5.45 | 20 | -46 | 6 |
| Cuneus | L | 0.001 | 5.07 | 4 | -84 | 28 |
| Cuneus | R | 0.051 | 6.14 | 8 | -80 | 28 |
| Lingual gyrus | R | 0.003 | 5.16 | 18 | -44 | -6 |
| Middle occipital gyrus | L | 0.012 | 6.58 | -52 | -72 | 16 |
| Middle occipital gyrus | R | 0.002 | 5.42 | 52 | -66 | 24 |
| Fusiform gyrus | L | 0.007 | 5.4 | -20 | -36 | -16 |
| Fusiform gyrus | R | 0.004 | 5.86 | 22 | -34 | -16 |
| Angular gyrus | L | 0.026 | 6.15 | -50 | -70 | 24 |
| Angular gyrus | R | 0.003 | 5.12 | 48 | -64 | 24 |
| PreCuneus | R | 0.003 | 5.51 | 20 | -46 | 8 |
| Middle temporal gyrus | L | 0.088 | 6.76 | -52 | -68 | 18 |
| Middle temporal gyrus | R | 0.095 | 7.74 | 52 | -68 | 14 |

### Hypothesis 3: Airborne Disease vs. Ancestral vs. Modern Fear Stimuli

To examine the neural response to airborne disease stimuli relative to ancestral and modern fear stimuli, paired t-tests were conducted for the following contrasts (see Figures 4b and 4c): (AF > AFsc) > (DIS > DISsc), (MF > MFsc) > (DIS > DISsc), (DIS > DISsc) > (AF > AFsc), and (DIS > DISsc) > (MF > MFsc). Whole-brain analyses were performed at p < 0.005, FWE-corrected. Detailed activation patterns for all contrasts are presented in Table 3, with key findings summarized below.

**Table 3.**
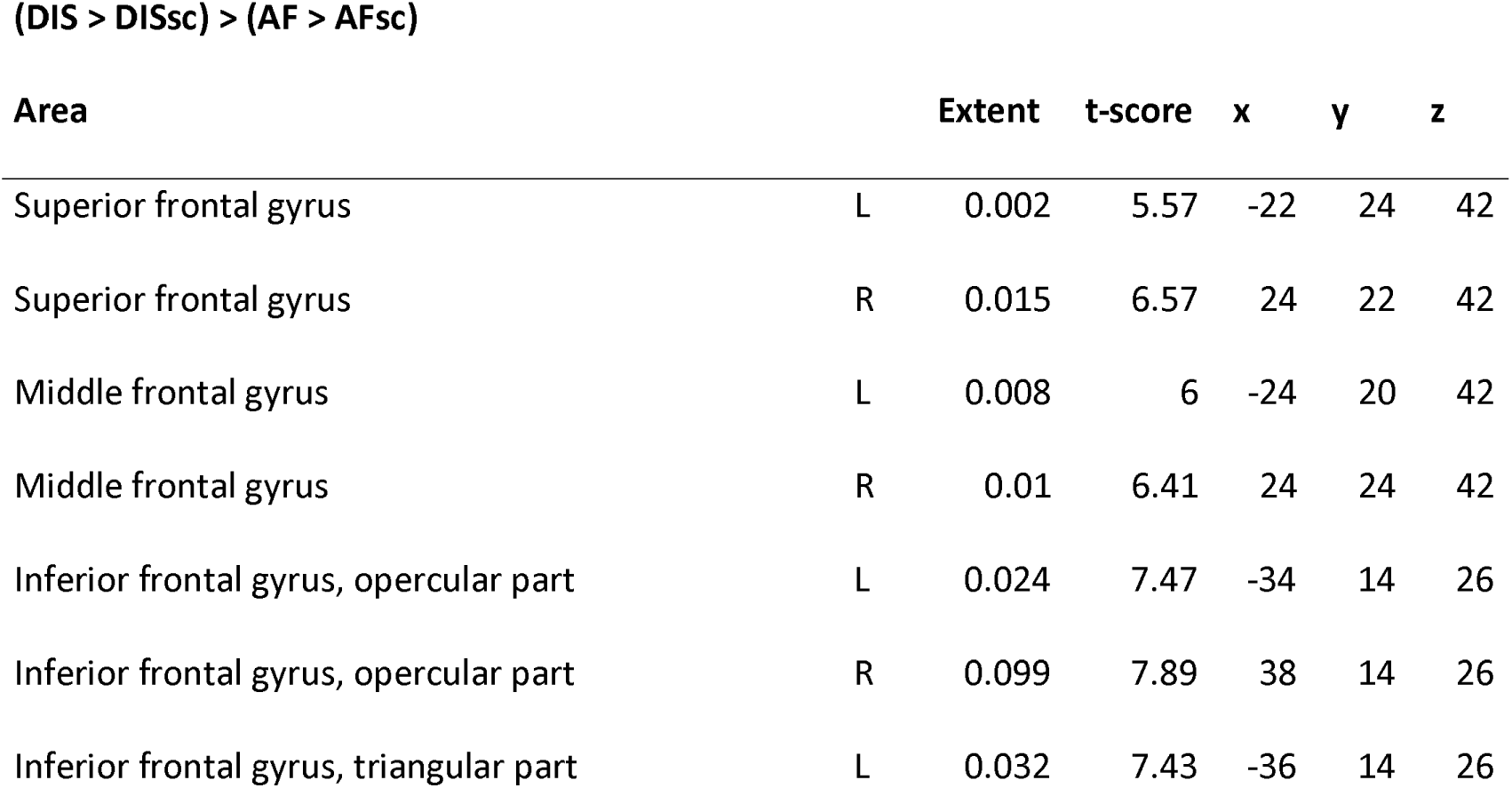
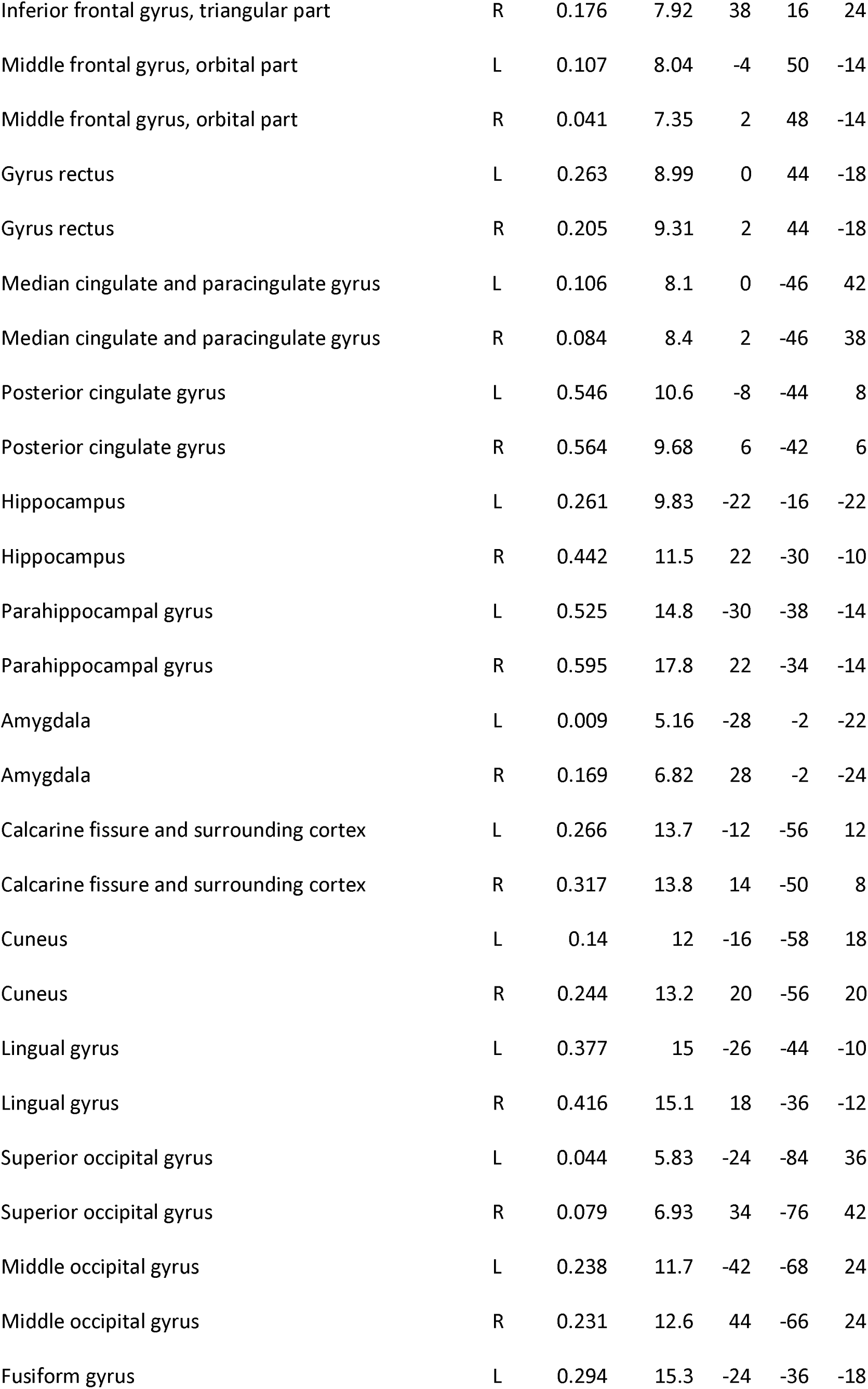
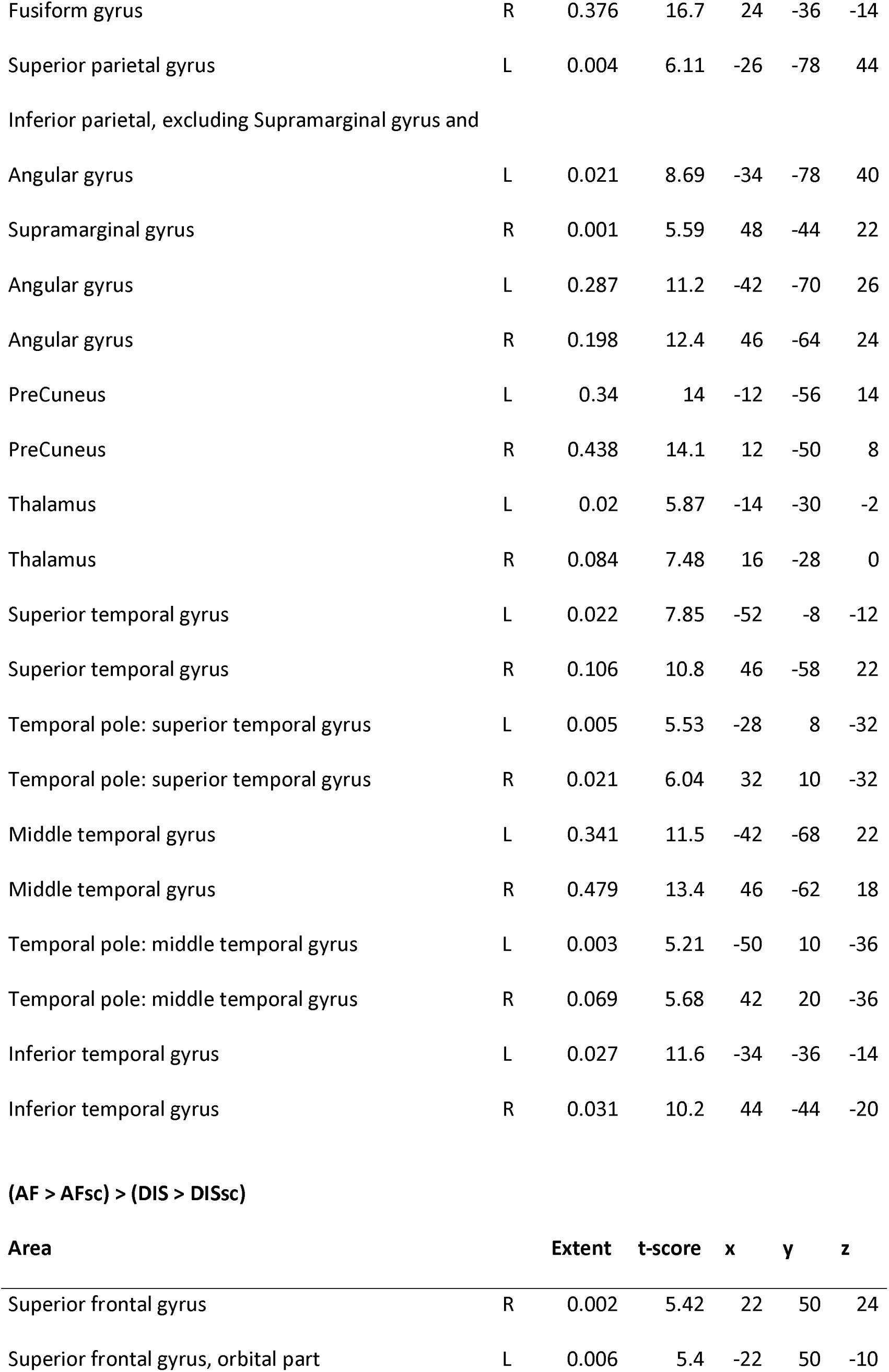
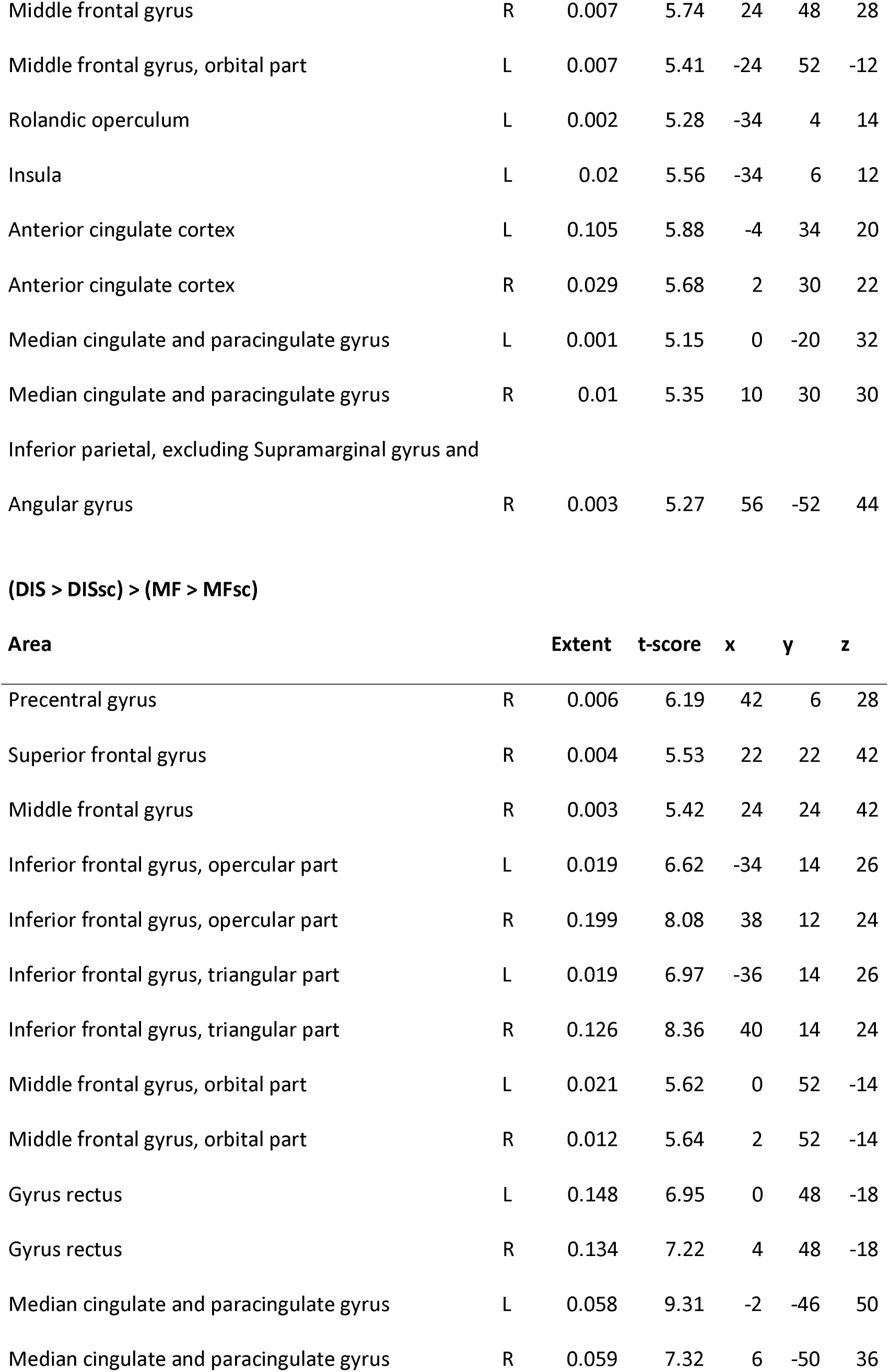
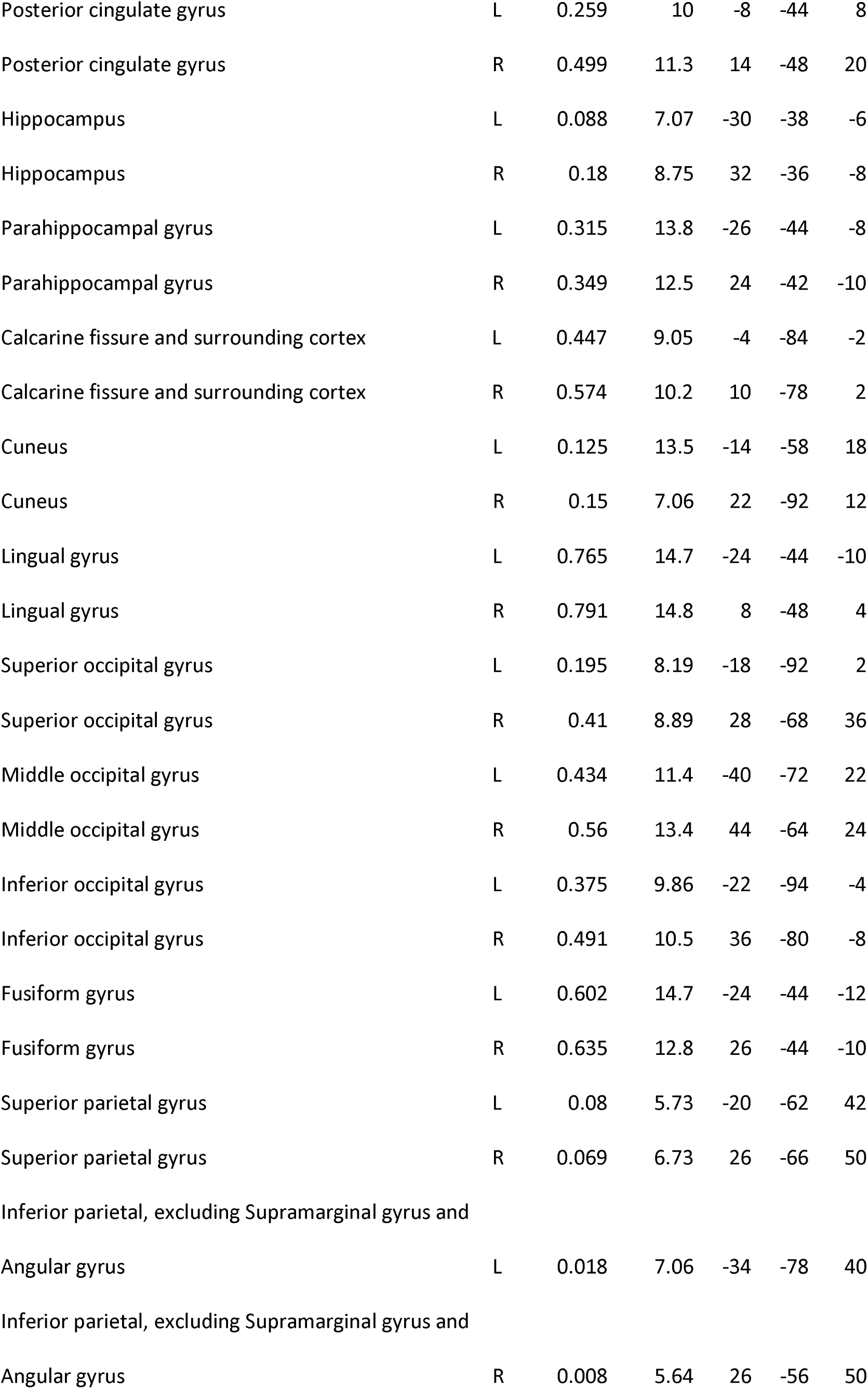
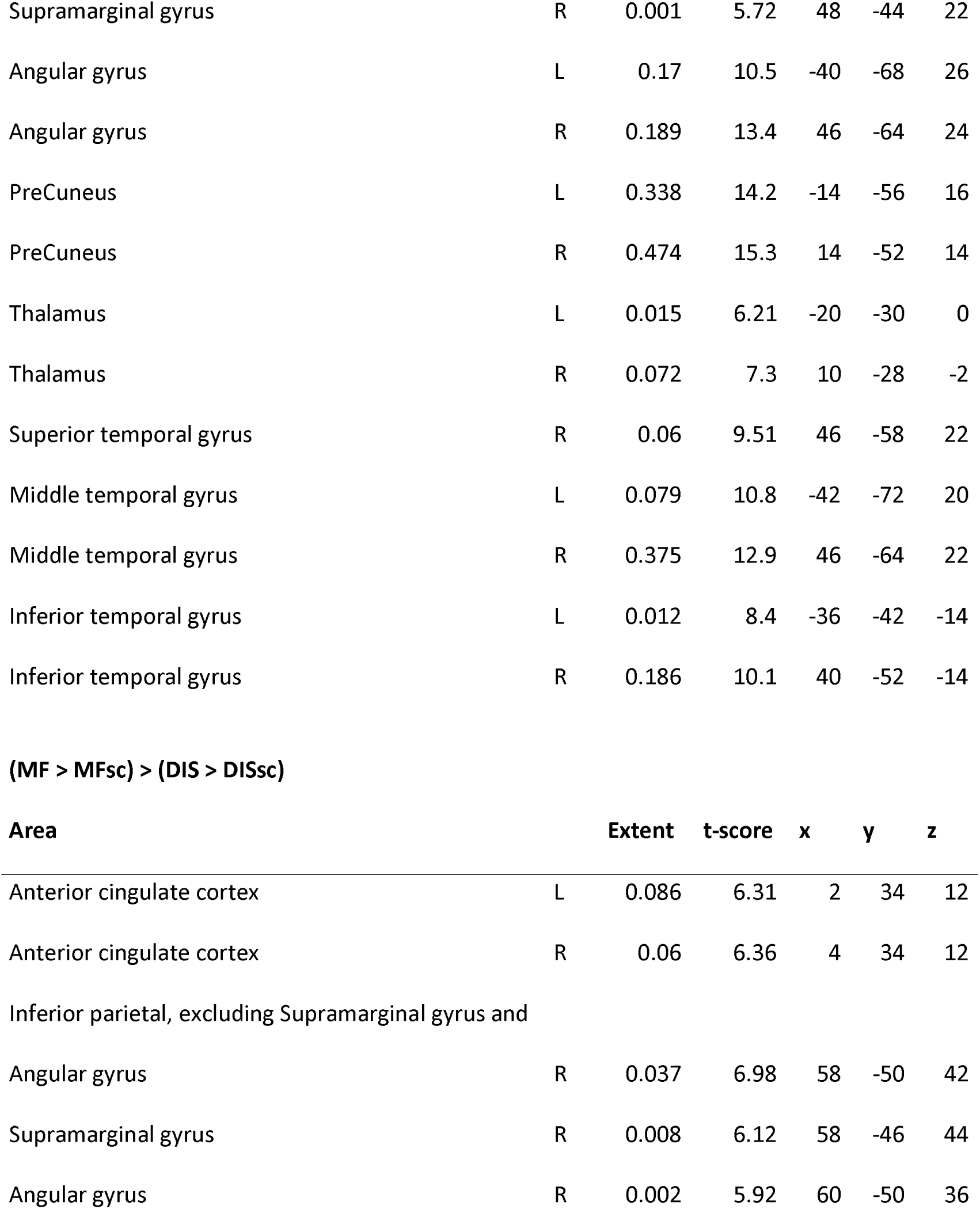
Contrasts (DIS > DISsc) > (AF > AFsc), (AF > AFsc) > (DIS > DISsc), (DIS > DISsc) > (MF > MFsc), and (MF > MFsc) > (DIS > DISsc); whole-brain analysis, thresholded at cluster level p<0.05 (FWE-corrected). The presented t-scores with the corresponding MNI coordinates (x,y,z) represent the maximum values within the brain area.”

### Ancestral Fear > Airborne Diseases

The contrast [(AF > AFsc) > (DIS > DISsc)] revealed heightened activation in regions associated with emotional processing and attentional engagement, including the frontal and parietal and limbic and cingulate areas, including bilateral anterior and median cingulate cortices, left insula, and left rolandic operculum).

### Modern Fear > Airborne Diseases

The contrast [(MF > MFsc) > (DIS > DISsc)] revealed significant activation in regions including the cingulate and parietal areas: bilateral anterior cingulate cortex and right inferior parietal gyrus, with additional activation in the right angular gyrus and supramarginal gyrus.

### Airborne Diseases > Ancestral and Modern Fear Stimuli

Significant activations were observed in a similar network of regions when compared to either ancestral [(DIS > DISsc) > (AF > AFsc)] or modern [(DIS > DISsc) > (MF > MFsc)] stimuli. The airborne disease stimuli elicited extensive activation across the frontal and limbic regions (bilateral superior, middle, and inferior frontal gyri, median and posterior cingulate cortices, bilateral hippocampus, and parahippocampal gyrus). Extensive activation was also found within the occipital and temporal areas (bilateral calcarine cortex, cuneus, lingual gyrus, fusiform gyrus, superior and middle temporal gyri, and inferior temporal gyrus), gyrus rectus (a part of the prefrontal cortex), angular gyrus and thalamus. These results suggest that monitoring stimuli depicting airborne diseases is cognitively demanding, requiring attention and prior experiences stored in episodic memory.

In addition, bilateral amygdala was activated during the contrast [(DIS > DISsc) > (AF > AFsc)]. Detailed activation patterns for all contrasts are provided in Table 3.

### Comparing the intensity and pattern of activation of the airborne diseases

As described in the Statistical analysis section, to examine the neural response to airborne disease stimuli relative to ancestral and modern fear, we computed the average activation index for three contrasts per subject (AF > AFsc, MF > MFsc, and DIS > DISsc). The corresponding distributions of indices over subjects are shown in Figure 5.

**Figure 5.** The Average activation index (AAI). Distributions of the AAI over subjects per condition. AF, MF, and DIS denote ‘ancestral fear’, ‘modern fear’, and ‘airborne disease’.

The activation index values were as follows: modern fear ranged from -6495 to 22240, with a mean of 7499; ancestral fear ranged from -1993 to 27374, with a mean of 12112; and airborne disease ranged from -2600 to 24188, with a mean of 15097. Pairwise comparisons revealed significant differences between the conditions: the activation index for modern fear differed from ancestral fear (p < 0.001), modern fear differed from airborne disease (p < 0.001), and ancestral fear differed from airborne disease (p = 0.013).

Finally, we fitted a PLS model that distinguished the activation maps obtained for modern and ancestral fear and applied it to predict one of two classes (‘MF’ or ‘AF’) for the activation maps for the airborne disease. Leave-one-out classification accuracy on the training data (AF + MF) was 61% (p-value, i.e. the chance to reach this accuracy by random choice with a success rate of 0.5, is 0.005). When applied to the DIS condition, the model predicted that 70% of the maps belonged to the MF class (p-value 0.001). That means that the activations for modern and ancestral fear show a significant difference, and airborne disease activation maps are closer to the modern than to the ancestral fear activation patterns.

## Discussion

### Methodological Validity: Ancestral Versus Modern Threats

In this study, we focused on differences in neural activation during exposure to ancestral and modern stimuli, as well as their comparison with stimuli depicting airborne diseases. To ensure that we measured what we intended to, we carefully selected stimuli that had been previously evaluated as highly fear-inducing (Peléšková, Polák, Janovcová, Chomik, Sedláčková, Frynta & Landová, 2024; Landová, Polák, Janovcová, Štolhoferová, Peterková, Chomik & Frynta, 2025). The results of this experiment confirmed our selection was appropriate as participants subjectively rated both ancestral and modern stimuli as strongly fear-inducing (significantly more so than disgust-inducing). However, to ensure the right interpretation of results, one additional step is needed – unambiguous categorization of the stimuli, which is not always present in existing literature.

In their ERP study, Brown, El-Deredy & Blanchette (2010) found that evolutionary and modern threats led to increased P1 amplitude, and the effect was stronger for modern threats (but see Zhang & Guo, 2019). However, they used knives and syringes as examples of “modern threat” stimuli. Though modern, these stimuli are sharp and could evoke ancestral fear responses due to their similarity to sharp objects encountered by early humans.

Similarly, Zsido, Csatho, Matuz, Stecina, Arato, Inhof, and Darnai (2019; see also Lipp, Derakshan, Waters, & Logies, 2004) used cats as controls in attention studies on snake fear. However, this approach overlooked the possibility that cats themselves could evoke fear responses due to their similarity to large felids, which are evolutionarily relevant predators or competitors (Frynta et al., 2023). Alternatively, cats might simply attract attention because of their high physical attractiveness to human observers (Landová, Poláková, Rádlová, Janovcová, Bobek & Frynta, 2018). Using ambiguous stimuli in studies comparing phylogenetic and ontogenetic threats can easily lead to biased results, as some authors might have (accidentally) found. It is quite possible that authors using ambiguous stimuli found the (expected) reaction towards ancestral stimuli, but wrongly interpreted the results as increased reaction towards modern threats (Shapouri & Martin, 2022).

In our study, we selected snakes as ancestral fear-inducing stimuli due to their well-documented status as ancestral threats, a topic that has been extensively studied and validated (Öhman & Mineka, 2003; Soares, 2010; Kawai, 2019). For modern stimuli, we deliberately avoided any that might resemble ancestral threats. Instead, we chose guns. Their shape does not correspond to anything humans would have encountered in the past, and their recognition as a threat is something learned during an individual’s lifetime.

A frequent methodological challenge in fMRI studies is ensuring the “comparability” of stimuli. When participants view an image, it affects them in multiple ways, not just emotionally (and each stimulus can evoke more than one emotion). Participants perceive elements such as color, light intensity, and the “comprehensibility” of the scene depicted, as they must interpret the image to some degree. When interpreting fMRI data, it is crucial to account for all these components, determine whether they are comparable across stimuli, and isolate the aspect of interest to the experiment. To address this issue, we compared each image first with its scrambled version. By “subtracting” the scrambled image response, we obtained a „purified“ reaction that included both the emotion evoked by the stimulus and its cognitive processing (mental interpretation and understanding of the image’s content). This approach filtered out the purely perceptual component. However, even with this method, we must consider the cognitive processing of stimuli in our interpretation. This processing is a critical aspect of responding to fear-inducing stimuli, encompassing not only the emotional fear response but also the assessment of the object’s level of danger, its distance, immediacy, and other factors.

First, in our study, we examined whether and to what extent the selected stimuli elicited activation corresponding to an emotional response (and potentially a response associated with emotional processing). Results showed that all three types of stimuli (ancestral fear, modern fear, and airborne diseases) activated extensive regions associated with the fear network and emotional processing of visual stimuli. The key region involved was the amygdala, which was bilaterally activated for all three conditions, along with the cuneus, lingual gyrus, fusiform gyrus, extensive occipital areas, and others (see the Results section). Additionally, regions processing attention and emotionally charged memories or contexts were activated, including the precuneus, thalamus, hippocampus, parahippocampal gyrus, and angular gyrus (Ranganath & Ritchey, 2012; Phelps, 2012). These findings suggest that our methodological approach was successful and that each of the stimulus blocks elicited the expected emotional response.

Of particular interest was the activation of the precentral gyrus and the basal ganglia, specifically the putamen and pallidum, in response to ancestral fear and airborne disease stimuli. These areas are responsible for the execution and regulation of involuntary and voluntary movements (Hassler & Dieckmann, 1969; Parkinson, McDonagh & Vidyasagar, 2009; Gerfen & Surmeier, 2011). During fMRI, participants are explicitly instructed to remain still to ensure valid data collection. However, intense fear can trigger bodily movements, such as trembling. Highly motivated participants likely suppressed these movements due to the immediate inhibitory action of the pallidum. The activation of the precentral gyrus in these two conditions supports this interpretation.

These results indicate that participants experienced more intense fear during the presentation of ancestral stimuli and airborne disease stimuli compared to modern stimuli, where such activation was absent. The middle frontal gyrus was also activated in both conditions, a region involved in the regulation of emotional responses through its connections with the limbic system (Ochsner et al., 2004). This further supports the notion of an intense emotional reaction. Additionally, the insula, part of the ventral attention network (or salience network), was activated during the ancestral threat (see below).

In contrast, modern stimuli (guns) activated additional regions, similar to airborne disease stimuli, including the superior and inferior frontal gyri (part of the dorsolateral and ventrolateral prefrontal cortex, dlPFC and vlPFC). These areas are involved in self-referential processing and introspection (Northoff, Heinzel, De Greck, Bermpohl, Dobrowolny, Panksepp, 2006), maintenance of working memory (Cabeza & Nyberg, 2000), and regulation of emotions and cognitive control (Aron, Robbins & Poldrack, 2004; reviewed in Webler, 2021). This suggests that modern stimuli, like airborne diseases, are more cognitively demanding to process. Evaluating these stimuli requires memory retrieval and an assessment of how dangerous the situation is in the observer’s context (self-reference). This interpretation is further supported by the increased activation of the gyrus rectus, which is primarily responsible for decision-making and social cognition (Hare, Camerer & Rangel, 2009).

A unique activation observed in response to airborne disease stimuli occurred in the medial cingulate cortex and the posterior cingulate gyrus, which are responsible for the cognitive assessment of imminent danger and cognitive evaluation of the situation (Fiddick, 2011; Pearson, Heilbronner, Barack, Hayden & Platt, 2011). These findings indicate that participants first assessed the airborne disease stimuli as not posing an immediate threat and then proceeded to evaluate the situation in greater detail. In contrast, encounters with a snake or a gun may be immediately perceived as acute dangers, leaving less time for complex situational analysis.

### Ancestral and Modern Threats Compared to Airborne Diseases

When comparing the individual contrasts, we consistently observed activation in regions associated with fear processing, such as the occipital areas, lingual gyrus, fusiform gyrus, and cuneus. These regions were activated in all pairwise comparisons: ancestral versus modern fear, modern versus ancestral fear, and airborne diseases compared to both fear-inducing conditions. Amygdala activation, however, was only observed in the contrast between airborne diseases and ancestral fear. This aligns with its relatively uniform activation across all affective stimuli, as statistical contrasts only reveal differences, not shared activations.

The most intriguing aspect of our findings lies in the distinct differences observed between the conditions. When comparing both ancestral and modern fear stimuli with airborne diseases, we found clear bilateral activation in the anterior cingulate cortex (ACC), a key node of the ventral attention network (Seeley et al., 2007; Kamran et al., 2014). The dorsal ACC is well-documented for its role in evaluating the salience of emotional stimuli (Etkin, Egner & Kalisch, 2011). Its activation in this context suggests engagement in assessing the immediate relevance and emotional intensity of these threats.

In comparison, the superior, middle, and inferior frontal gyri, along with the orbital part of the middle frontal gyrus and gyrus rectus, were activated when watching the airborne diseases (when directly compared to the ancestral and modern fear). These regions are more prominently involved in cognitive regulation of emotions, decision-making, and attention control. They are often recruited when individuals process complex or abstract threats (Wager, Davidson, Hughes, Lindquist & Ochsner, 2008; Ochsner, Silvers & Buhle, 2012). Their activation during airborne disease stimuli suggests that these regions were engaged for cognitive evaluation, such as integrating epidemiological knowledge, symbolic cues (e.g., masks, hospitals), and longer-term implications of the threat. Unlike the ACC, these areas are less involved in immediate threat detection and more in deliberative processes, such as developing behavioral strategies (e.g., deciding to wear a mask or avoid crowded areas).

These results align with additional analyses comparing the overall similarity and intensity of activation patterns for airborne diseases with those elicited by ancestral or modern stimuli. When directly comparing contrasts, areas associated with complex cognitive threat evaluation appeared prominently only for airborne diseases, likely due to their heightened intensity. However, analyses comparing stimuli with scrambled controls revealed that these regions also activate during the processing of guns, as confirmed by the partial least squares (PLS) analysis. This analysis shows that the overall activation pattern—specifically, “which regions” are activated—resembles that of modern threats for airborne diseases. In contrast, the activation intensity, as indicated by both contrast analyses and average activation index (AAI) measures, is more akin to ancestral threats.

### Neural Activation and Subjective Ratings

In previous studies comparing airborne diseases with ancestral and modern stimuli (Peléšková et al., 2024; Landová et al., 2024), participants rated stimuli based on fear, disgust, and anger they experienced. Ancestral and modern stimuli, as well as those depicting airborne diseases, were further categorized into primarily fear- or disgust-inducing subcategories. Results showed that modern threats (car accidents, electricity) elicited the highest fear ratings, followed by ancestral threats (snakes, heights). Airborne diseases (pandemic fear, described here through scenarios involving people wearing masks and overcrowded hospitals), categorized as intermediate between ancestral and modern threats, were rated as eliciting the least fear. In contrast, ancestral disgust stimuli (body waste products, worms) and “pandemic disgust” (coughing and sneezing individuals) elicited the highest disgust ratings, with the latter overlapping substantially with ancestral disgust stimuli (Landová et al., 2024).

In the present study, airborne disease stimuli corresponded to “pandemic fear” scenarios described in the two previous studies (i.e., masked individuals, crowded hospitals). Results differed, with ancestral stimuli and airborne diseases evoking stronger neural responses compared to modern stimuli. Several factors could explain this discrepancy: First, subjective evaluations may not fully correspond to neural processing. Conscious ratings might reflect the outcome of emotional regulation rather than raw neural responses (Andrews-Hanna, Smallwood, & Spreng, 2014; Haspert, Wieser, Pauli, & Reicherts, 2020). Neural responses captured by fMRI, such as amygdala activation, could represent immediate, unregulated emotional reactions, whereas subjective ratings involve reflective appraisal.

Another reason could be differences in processing modalities. Reading about ancestral threats may not elicit the same neural responses as direct exposure, as these threats are typically processed through fast, automatic pathways within the fear network. Reading text, however, engages cognitively demanding processes requiring attention and comprehension, activating regions such as the left inferior frontal gyrus, bilateral dorsolateral prefrontal cortex (middle and superior frontal gyri), and bilateral hippocampus (Keller, Mason, Legg, & Just, 2024). In our study, these regions were predominantly activated during the processing of modern stimuli and airborne disease scenarios. But in textual form, the distinction between ancestral and modern stimuli can be blurred if all stimuli need to be processed by these regions first.

Third, the cultural context and stimulus familiarity might have played a role. In contrast to Peléšková et al. (2024) and Landová et al. (2024), where modern stimuli included scenarios like exposed electricity and car accidents, this study used guns. In the Czech Republic, guns are less commonly experienced in threatening contexts and could be more likely associated with hunting or other neutral-to-positive activities. In contrast, many participants may have direct, impactful experiences with electricity or car accidents, which could make these stimuli more salient and emotionally evocative in the prior studies.

These findings are consistent with Dhum et al. (2017), who reported stronger subjective ratings for modern threatening stimuli but stronger neural activation in fear-processing networks, including bilateral amygdala activation, for ancestral stimuli. The authors argued that while subjective ratings may involve complex cognitive evaluations, they do not always align with amygdala activation, which corresponds more closely to the intensity of emotional valence (Anders, Eippert, Weiskopf, & Veit, 2008).

In our study, amygdala activation was observed across all stimuli, with the most pronounced response in the contrast airborne diseases > ancestral fear. Similarly, this pattern does not align with subjective fear ratings, because both modern and ancestral stimuli were rated as more fear-inducing than airborne diseases. These discrepancies underscore the complex interplay between subjective experiences and neural responses, highlighting the need for integrated approaches that account for both dimensions.

### Context of Human Presence

Studies by Yang and colleagues (Cao, Zhao, Tan, Chen, Ning, Zhan, & Yang, 2014; Fang, Li, Chen, & Yang, 2016) explored how the amygdala reacts to negative (and neutral) stimuli involving animals and objects depending on the context of human presence. Both studies consistently found that in images without humans, the amygdala showed a stronger response to negative animals. However, when human presence was included in the context, the results diverged: Fang et al. (2016), in the case of subliminal presentation, reported even stronger amygdala activation, whereas Cao et al. (2014), using supraliminal (conscious) presentation, observed greater amygdala activation for negative objects, with additional top-down modulation by the anterior prefrontal cortex.

In our study, animals were depicted only in the absence of humans. Negative modern threats, however, were exclusively presented within a human context, as these are more intense in such settings (especially with guns; Yang, Wang, Yan, Zhu, Chen, & Wang, 2012). This likely explains why, similar to the stimuli depicting airborne diseases, we observed activation in the vlPFC (inferior frontal gyrus) and vmPFC (gyrus rectus). These areas are implicated in inhibitory control and the regulation of responses during social interactions (Smith, Nelson, Rappaport, Pine, Leibenluft & Jarcho, 2018; Nelson & Guyer, 2011), as well as in integrating social values and personal relevance, mediating empathy and emotional responses in social contexts (Winecoff, Clithero, Carter, Bergman, Wang & Huettel, 2013; Hiser & Koenigs, 2018). The amygdala, however, was activated across all conditions.

Nonetheless, our study differs substantially from Yang’s work as their focus was on negative valence rather than fear, and their stimuli (e.g., “ugly” beetles) reflect this distinction. Consequently, their findings are not directly applicable to ancestral threats, as suggested in reviews like Shapouri & Martin (2022), because stimuli with negative valence are not necessarily life-threatening.

## Conclusions and practical implications

This study demonstrates that the human brain exhibits heightened sensitivity to ancestral threats when compared to modern threats. Specifically, the basal ganglia and ventral attention network (ACC, insula) show stronger responses to stimuli representing ancestral threats. These findings underscore the enduring influence of evolutionary pressures on neural processing, suggesting that certain brain circuits have retained their sensitivity to survival-relevant stimuli from our evolutionary past.

Moreover, the results reveal that while ancestral threats elicit stronger and broader neural responses than modern threats, the neural response to airborne threats demonstrates an intriguing duality. By intensity, the brain’s reaction to airborne threats mirrors its heightened sensitivity to ancestral dangers. However, the patterns of neural activation more closely resemble those observed for modern threats, with predominant activation in cortical regions associated with analytical evaluation and context-specific decision-making. These findings suggest that while the human brain retains an evolutionary bias favoring intense responses to survival-relevant stimuli, it also recruits specialized circuits for processing airborne threats, likely reflecting their abstract, socially mediated nature.

From a practical standpoint, these insights have critical implications for addressing future pandemics. Understanding how the brain processes these threats can inform mental health interventions targeting pandemic-related anxiety and stress, particularly by addressing the dual demands of immediate threat perception and sustained cognitive vigilance. Future research should explore how neural processing of airborne threats evolves over prolonged exposure, offering further strategies to bolster public resilience during global health crises.

## Data availability statement

The original contributions presented in the study are included in the article/Supplementary material, further inquiries can be directed to the corresponding author.

## Acknowledgements

We would like to thank all our respondents who helped us with their kind participation to conduct this study. We would like to thank Associate Professor Jaroslav Tintěra for his help in programming the fMRI experiment and Kristýna Sedláčková for help with recruitment of the participants. We also thank Dr. Filip Španiel for inspiring discussions regarding the design of the experiment and for providing inspiring insights into the nature of the human brain. We thank Rudolf Gašpar and Jiřina Golovinská for their help with the administration of the accompanying tasks.

## Author contributions

EL, DF, SR, and JH — conceived and designed the research; SR — prepared the stimuli, AC — recruited the participants and collected the data; AP and DT — curated the data; DT and AP — fMRI data visualization, DT, AP, and JH — analyzed the data; SR, AP, and EL — wrote the first draft of the manuscript; EL, DF, and JD — reviewed and edited the manuscript; EL — acquired the funding. All authors approved the final version of the manuscript.

## Supporting information captions

**S1 Supplement.** List of sources of the photos used in the study. Some of the pictures from Pixabay and other sources were altered, cropped, or color-adjusted. The pictures from affective online databases were unaltered.

**S2 Supplement A.** Contrast ancestral fear > scrambled ancestral fear (AF > Afsc), whole-brain analysis, thresholded at cluster level p<0.05 (FWE-corrected). The presented t-scores with the corresponding MNI coordinates (x,y,z) represent the maximum values within the brain area.

**S2 Supplement B.** Contrast modern fear > scrambled modern fear (MF > MFsc), whole-brain analysis, thresholded at cluster level p<0.05 (FWE-corrected). The presented t-scores with the corresponding MNI coordinates (x,y,z) represent the maximum values within the brain area.

**S2 Supplement C.** Contrast airborne disease > scrambled modern disease (DIS > DISsc), whole-brain analysis, thresholded at cluster level p<0.05 (FWE-corrected). The presented t-scores with the corresponding MNI coordinates (x,y,z) represent the maximum values within the brain area.

